# Habitual digital media use and the brain: a meta-analysis

**DOI:** 10.64898/2026.03.05.709910

**Authors:** Lena J. Skalaban, Ashley A. Murray, Jason M. Chein

## Abstract

Research on the relationship between digital media and neurocognitive function has blossomed with the rising digital age and advent of social media, producing a growing literature focused on how technological developments may be affecting users’ brains. Much of the science has focused on the involvement of specific brain systems that support reward (e.g., nucleus accumbens, orbitofrontal cortex), cognitive control (e.g., lateral prefrontal, anterior cingulate), and socio-emotional processes (e.g., temporo-parietal junction) and why they might be especially relevant to digital media engagement. However, a broad and systematic analysis of the consistency of findings across neuroimaging studies has not yet been published. Here, we conducted a coordinate-based meta-analysis based on published structural and functional MRI studies exploring habitual digital media engagement. Adopting a granular approach to summation of this literature, we use Activation Likelihood Estimation (ALE) and find that the most consistent effects arise in the anterior insular cortex, a region implicated in the integration of social and emotional information that has not been frequently highlighted in the prior literature on digital media effects in the brain. This discovery encourages reconsideration of how the brain is likely to affect, and be affected by, digital media engagement and online behavior.

## INTRODUCTION

Digital media engagement in the general population has increased significantly over the last two decades (Korte, 2020), with over 90% of all American adults now owning a smartphone (Pew Research, 2024) and approximately 310 million Americans (93% of the population) active on social media (Statista, 2025). This trend has greatly impacted how humans socialize, learn, and spend their recreational time. This consequential change in daily behaviors has also spurred research into the potential impacts that digital media and technology use, especially smartphone and social media use, might be having on neurocognitive function –– with the majority of papers on this topic published in the last 10 years.

While work in this area is often dominated by concern regarding the potential effects of digital media habits on behavior and mental health (Keles et al., 2020; Lissak et al., 2018; Ra et al., 2018), there is a complementary and growing body of work exploring how digital media and technology use habits may also be reflected in brain structure and function (Chiu & Chein, 2022; Choudhury & McKinney, 2013; Hutton et al., 2024; Wadsley & Ihssen, 2023). Many studies exploring the relationship between brain and digital media behavior home in on specific brain areas and networks that are most likely to be implicated in digital media behaviors based on our understanding of their function. For example, studies typically probe the prefrontal cortex/frontoparietal control system (Brand et al., 2014; Sun et al., 2012); the anterior cingulate cortex/salience network (Schmitgen et al., 2020; Wang et al., 2017); the nucleus accumbens/reward circuity (Ko et al., 2009; Schmitgen et al., 2025); or socio-emotional processing regions like the amygdala (He et al., 2017; Zhao et al., 2025) for evidence of differentiation that is dependent on one’s history or intensity of digital media involvement.

Likewise, several excellent theoretical and systematic reviews of work in this research area (Chiu & Chein, 2022; Crone & Konijn, 2018; Dores et al., 2025; Firth et al., 2019; Uncapher & Wagner, 2018; Wadsley & Ihssen, 2023) are organized around a set of guiding assumptions for why brain systems supporting executive control, reward valuation, and social information processing might be inculcated in these digital media behaviors; and accordingly, these reviews highlight empirical findings that align with those assumptions. While this approach to summarizing the evidence has helped advance our understanding of this topic, a now enlarged empirical neuroimaging literature presents the opportunity to identify the key brain loci of digital media habits in a complementary, but more quantitative, matter that is agnostic to theoretical assumptions.

To this end, the current study pairs qualitative summary with a quantitative approach that interrogates the spatial consistency of the neuroimaging findings surrounding digital media use habits. Specifically, we deploy coordinate-based meta-analysis (CBMA) methods to form Activation Likelihood Estimates (ALEs; Turkeltaub et al., 2002; Wager et al., 2003), adopting broad and liberal study inclusion criteria to investigate where group and individual differences in digital media engagement most consistently coincide with variation in brain function and structure. CBMA allows us to collate findings that span different analysis methods, varied sample sizes, and distinct subject populations. Importantly, this method is also data driven, and thus, provides an objective method for detecting consistent effects. As such, it can reveal consistencies in brain areas that may be under-emphasized in work guided by *a priori* theoretical frames and can test whether the brain systems most commonly emphasized in individual empirical papers and reviews actually warrant such emphasis.

## METHODS

### Data search strategy

We adopted an encompassing approach that included data from studies using structural and functional MRI, conducted in both developing (e.g., adolescent) and adult populations, and measuring varied forms of digital media usage. In accordance with PRISMA reporting guidelines, Figure 1 provides a summary of our search and paper selection strategy. We collected papers from multiple databases (Google Scholar and PubMed) and cross-referenced them. We used the following search terms to identify papers that contained reference to some form of neuroimaging and any one of many terms that fall under the purview of “digital media” use: “(MRI OR neuroimaging) AND social media OR digital media OR screen time OR smartphone OR social networking”. Following the broad initial search, we eliminated studies that were not published in English (n = 2) and removed any duplicates that resulted from cross-referencing of multiple databases (n = 109). The initial search was conducted by all three authors and then any duplicates found by multiple authors were eliminated.

**Figure 1.**
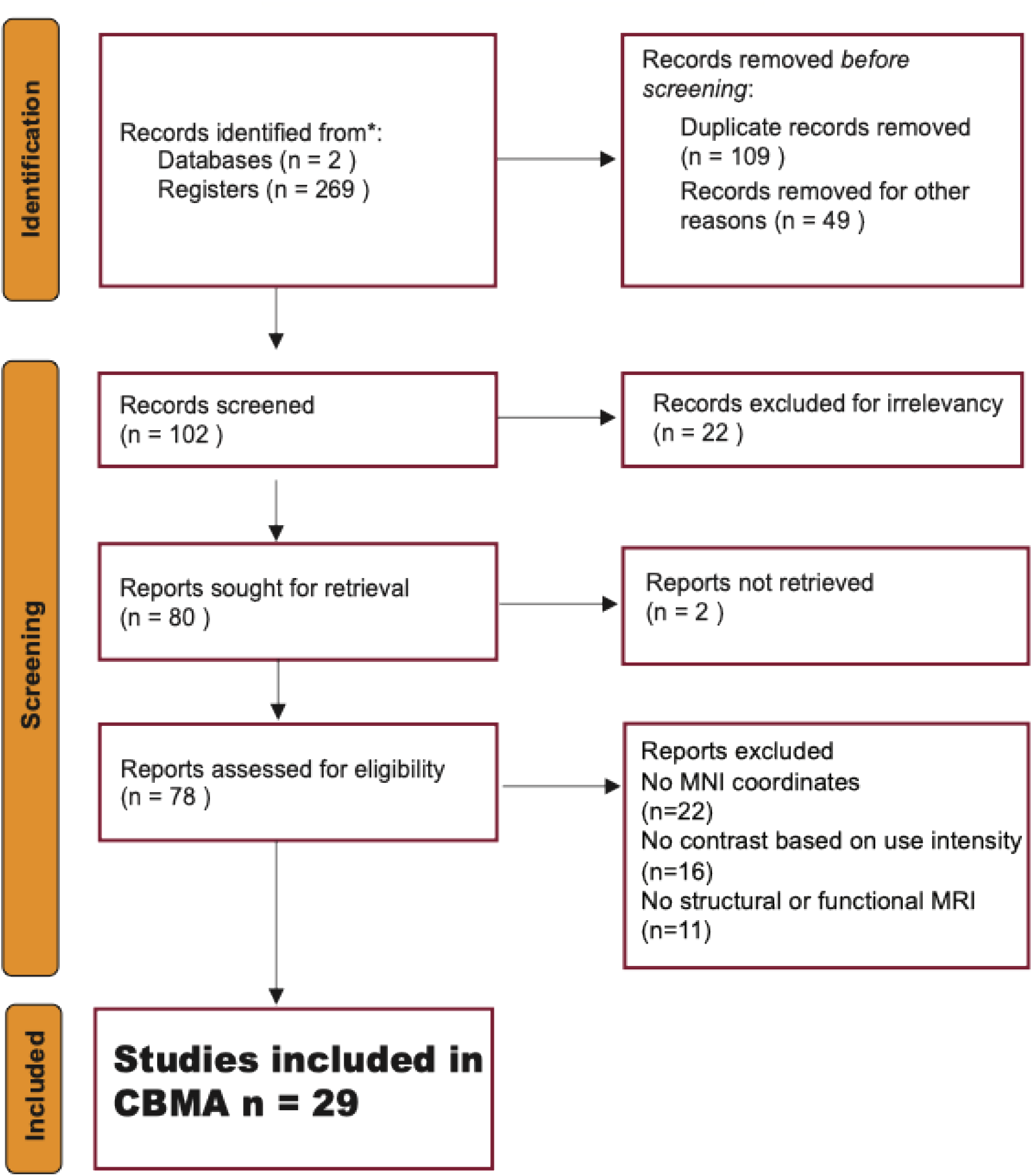
Prisma diagram summarizing study search, selection and inclusion. Studies included in the meta-analysis went through identification and screening processes. Records were accessed through two databases yielding 269 registers. After removal for irrelevancy, duplicates, non-English language publication, or inability to access the original manuscript, reports were assessed for eligibility. Reports were excluded if they did not contain stereotaxic f/MRI coordinates usable in an CBMA meta-analyses yielding 29 studies.

### Study selection

Following the search, first and second authors reviewed titles and abstracts for relevancy (n = 102). We restricted our analyses to studies focused specifically on modern forms of digital media usage (e.g., social media, smartphone screen-time, internet use habits) and excluded studies (n = 22) investigating niche behaviors or narrow patterns of addiction (e.g., “internet gambling”, “online shopping addiction”) or focused on traditional media forms (e.g. television, print). We also eliminated any articles that mentioned, but did not actually include, structural or functional MRI data and analyses (n = 11). The set was further restricted to studies that reported neuroimaging findings based on direct statistical comparisons between higher (e.g., heavy, excessive, problematic, addicted) versus lower (e.g., control) use groups, or measured individual differences in digital media habits/usage intensity in a continuous manner. This constraint excluded several task-based functional MRI studies designed to isolate the neural correlates of experimentally-manipulated facets of digital media behavior (e.g., responsivity to social media images with a manipulated number of “likes” - Sherman et al., 2018; Go/Nogo inhibition of responses to social-media-associated vs. neutral stimuli - Turel et al., 2014), but didn’t report statistical comparisons across groups or individuals who differed in their use habits (n = 16).

Exclusion of this latter type of study precluded potential biasing of CBMA findings simply due to the frequency with which a given process (e.g., responsivity to social media “likes”) is studied in the literature even when this process may not differentiate individuals who engage more heavily in the behavior from those who do not. Lastly, the purposes the CBMA, we also eliminated any study that did not report spatially normalized coordinates (n = 22). All studies that survived the inclusion criteria and reported coordinates for significant loci did so using Montreal Neurological Institute, MNI, convention). This resulted in 29 studies in total. Information on all papers included in the CBMA including citations can be found in Supplemental Table S1.

### Data extraction

Data was successfully extracted from all 29 studies that passed other inclusion criteria. That is, if any study included stereotaxic coordinates, without any further consideration of study quality, was included in an ALE meta-analysis. All articles were reviewed by at least two coders (first and second authors). Information extracted from each paper included citation, sample size, age of participants (when available), imaging modality (structural or functional MRI), media type measured (social media or other digital media), and directionality of reported effects (positive or negative). For structural MRI studies, ‘positive’ directionality meant that the study observed a relative increase in gray matter (e.g., cortical thickness, volume) among the relatively higher digital media use group/individuals. For functional MRI studies, ‘positive’ directionality meant greater activation and/or inter-regional connectivity for the higher use group/individuals. We also recorded the specific region labels used by each paper to identify significant ROIs or regions showing whole-brain effects. The first and second authors completed and validated data extraction of x, y and z MNI stereotaxic coordinates from each paper. Any article for which there were questions or discrepancies was reviewed by all three authors.

### Activation-likelihood estimation analyses

To conduct the CBMA, we adopted an activation likelihood estimation (ALE) approach (Turkeltaub et al., 2002; Chein et al., 2002) using the NiMARE (Neuroimaging Meta-Analysis Research Environment) meta-analysis package in Python 3 (RRID:SCR_017398; Salo et al., 2022) ALE analyses create thresholded brain maps identifying regions of greatest spatial consistency in the occurrence and location of the extracted coordinates. In ALE, each entered coordinate (typically representing the peak regional voxel for a relevant statistical contrast) is modeled as a three-dimensional gaussian distribution centered on that coordinate and adjusted in value as a function of study sample size. These coordinate-centered distributions are summed to produce a consensus map. The resulting consensus map is then compared to an empirical null distribution created via permutation simulations for the purpose of statistical thresholding. All ALE analyses were run with 10,000 permutations and were probability thresholded using Montecarlo family-wise error correction with a cluster defining p-value threshold of *p* =.05.

When this threshold produced large clusters that encompassed multiple functional regions, we repeated the analysis at a stricter cluster defining threshold of *p* =.001. Of note, ALE permutations are sensitive to both the count and distribution of the study coordinates that are entered into the analysis, and thus, specific subsets of studies can produce novel ALE clusters that do not surpass the correction threshold when a more inclusive set of studies are entered together. To obtain consensus region labeling for each resulting significance cluster, the peak x, y, and z coordinates from the cluster were submitted to Neurosynth (Yarkoni et al., 2011) and the highest probability anatomical label for each locus was identified. We reviewed the Neurosynth outputs in combination with visual inspection of ALE statistical maps to confirm accurate labeling for each peak region..

We generated several ALE maps, based on different groupings of the available studies. Any paper surviving the inclusion criteria was included in an Overall ALE map (n = 29). We also composed an ALE that separated out the subset of studies adopting a spatially unbiased whole-brain analysis (n = 19) from those that only interrogated restricted portions of the brain using a region-of-interest (ROI) and/or network-based approach (n = 10), which could have biased the localization of consensus regions. From the full set of 29 studies, we also constructed ALEs for the subset of studies using only structural MRI, using only functional MRI, and those studies focusing uniquely on use of social media (i.e. social networking sites). Of note, while most of the selected studies relied on adult participant samples, some studies were conducted in children and adolescents (information on which studies included developmental samples is provided in Supplemental Table S1). While we had planned to conduct separate ALEs for adults and developmental populations, only 8 of 29 selected studies included developmental samples (and these were not always separated out from the adult sample). Thus, there were too few studies across too wide an age range to support separate ALEs for developmental age groups. The meta-analysis methods for this study were not pre-registered.

### Focus count-analysis

Focus-count analyses were conducted on both the overall ALE and each of the subset ALEs. These analyses detail the relative contribution of individual studies to any significant cluster identified in an ALE map. As such, these analyses estimate the number of datapoints/studies contributing to a significant ALE cluster and can also be used to flag papers that have an especially large influence on a computed cluster. We used these focus-count tables to determine the consistency of effects, and to assess how often effects contributing to a specific consensus region were in the positive (i.e. stronger activation/increased gray matter for heavier digital media users) or negative (e.g. weaker activation/reduced gray matter for heavier digital media users) direction.

### Studies meeting criteria but not reporting coordinates

In addition to the papers included in the ALE analyses, our search produced 22 papers that fulfilled all study selection criteria but reported significant findings only by reference to functionally or anatomically pre-defined regions, and without reporting stereotaxic coordinates for observed effects. Papers that fell into this category comprised studies using *a-priori* ROI analyses (e.g., based on atlas-defined regions) and studies reporting analyses summarizing patterns observed across large-scale networks of regions (e.g., frontoparietal network, default mode network). While these papers could not be added to the CBMA (because there were no coordinates to extract), they detail findings that can be considered in relation to the CBMA results and that have influenced current beliefs about how and where digital media use habits manifest in the brain. We therefore tabulated observations from these studies (see Supplemental Table S3) and consider how the pattern of overall findings aligns with findings emerging from the current ALEs.

## RESULTS

### ALE analyses

*Overall ALE analysis* (all 29 studies). We first ran an ALE analysis based on the complete sample of 29 functional and structural MRI studies (30 contrasts) identified through our screening process. This ALE revealed two significant clusters, one centered in the right anterior insula (x = 30, y = 20, z =-8; Figure 2A) and the other (relatively weaker) cluster with its peak in the precuneus (x = 6, y = −68, z = 50; Figure 2B; Table 1). The larger anterior insula cluster extended laterally into the opercular inferior frontal gyrus and medially into the vicinity of the ventral striatum. At a stricter cluster defining threshold of p <.001, only a highly circumscribed anterior insula peak survived (see Supplemental Figure S1A).

**Table 1.** Peaks and Focus Counts from each ALE. Focus count (FC) indicates the number of studies that contributed to the cluster; +:− shows the ratio of the number of studies reporting positive (i.e. greater activity, greater volume/thickness) vs. negative (weaker activity, reduced volume/thickness) effects for heavier digital media users. Note that sometimes effects occurred in opposing directions for different conditions within a single study. AIC = Anterior Insular Cortex, IFG = Inferior Frontal Gyrus.

| ALE | #Studies | Right AIC+IFG |  |  |  |  | Left AIC+IFG |  |  |  |  | Precuneus |  |  |  |  |
| --- | --- | --- | --- | --- | --- | --- | --- | --- | --- | --- | --- | --- | --- | --- | --- | --- |
|  |  | *x* | *y* | *z* | FC | +/- | *x* | *y* | *z* | FC | +/- | *x* | *y* | *z* | FC | +/- |
| OVERALL | 29 | 30 | 20 | -8 | 13 | 5/9 |  |  |  |  |  | 6 | -68 | 50 | 5 | 2/3 |
| Whole-Brain | 19 | 26 | 16 | -8 | 9 | 5/5 | -34 | 18 | -4 | 5 | 4/2 |  |  |  |  |  |
| Functional | 20 | 26 | 22 | -4 | 8 | 5/4 | -36 | 16 | -4 | 8 | 4/5 | 6 | -68 | 48 | 5 | 2/5 |
| Structural | 9 | 32 | 16 | -10 | 4 | 0/4 |  |  |  |  |  |  |  |  |  |  |
| Social Media | 10 | 24 | 12 | -8 | 5 | 2/4 |  |  |  |  |  |  |  |  |  |  |

**Figure 2.**
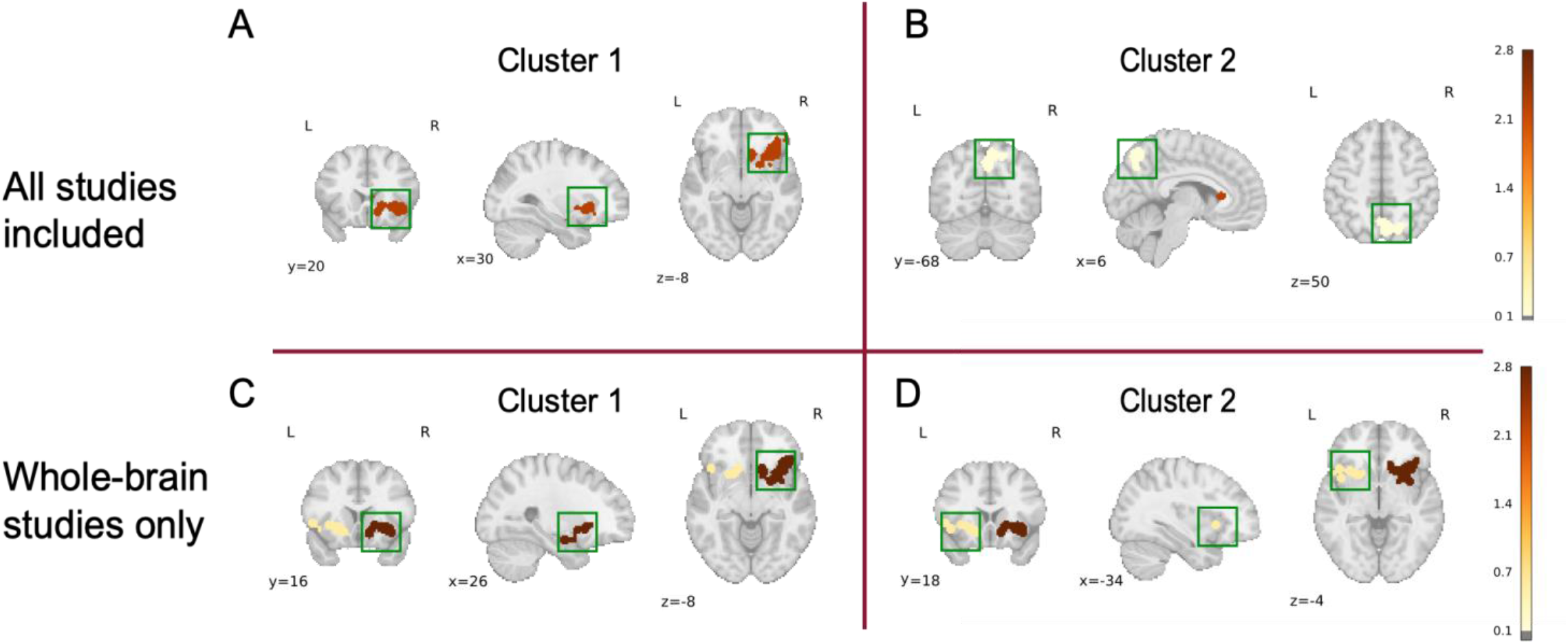
Overall ALE and whole-brain only ALE. Two ALEs were run, one with all studies included, and one with only studies reporting coordinates from whole-brain analyses. The overall ALE revealed two supra-threshold clusters. Cluster 1 was approximately centered in the right anterior insula, while Cluster 2 was centered approximately in the precuneus. The whole-brain only ALE also produced two clusters; Cluster one centered approximately in the right anterior insula and cluster two centered around the left anterior insula, with both clusters extending toward the midline.

A focus-count analysis for the regions surviving correction in this map revealed that the anterior insula cluster (Cluster 1) was supported by 13 of 29 studies, demonstrating moderate consistency, while the cluster centered in the precuneus (Cluster 2) had contributions from only 5 of 29 studies, indicating a considerably less robust and consistent pattern (see Supplemental Table S2 for full report of the focus counts). Specific studies also made an outsized contribution to each of the anterior insula (Maza et al., 2023, n = 178) and precuneus (Henemann et al., 2023, n = 132) sites, due to their relatively large sample sizes and coordinate proximity to the cluster center.

*Whole-brain only ALE* (19 studies). The full corpus of studies meeting eligibility criteria for the overall ALE analysis included 19 whole-brain studies and 10 studies that relied on regionally-constrained (ROI, network-based) approaches. In principle, the inclusion of the regionally-constrained studies could have biased the overall ALE to favor regions of *a priori* theoretical interest (of the relevant studies in the overall sample, 5/29 included an ROI specifically in anterior insula and 4/29 included an ROI specifically in the precuneus). To address this potential concern, we also computed a more restricted ALE that included only coordinates arising from the 19 studies using whole-brain contrasts, allowing us to compare the impact of including, or not including, coordinates from ROI- and network-based contrasts. This whole-brain only ALE produced two significant clusters: one cluster was again centered in the right anterior insula (Figure 2C), with a peak coordinate (x = 26, y = 16, z = −8) and extent that mirrored the right anterior insula cluster produced by the overall ALE analysis. The second cluster identified in this whole-brain ALE was centered in a homologous anterior insula region in the left hemisphere (x = −34, y = 18, z = −4; Figure 2D; Table 1; see also Supplemental Table S3B) that had a size and shape similar to its right-lateralized counterpart. Using a focus-count analysis, we determined that the first cluster demonstrated relatively high consistency (9 of 19 studies), while the second showed less cross-study consistency (5 of 19 studies).

To characterize potential differences across studies using different imaging modalities (functional vs. structural), and to focus in on a more homogenous form of digital media behavior (social media), we conducted a series of additional targeted ALE analyses. Specifically, we coded studies according to whether they used functional imaging (n = 20) or structural imaging techniques (n = 9), and based on whether they assessed variation in social networking site use (n = 10) or relied on other, more heterogenous, indices of habitual digital media behavior (n = 19). ALE findings from the heterogenous category of “other digital media behavior” can be found in supplemental analyses.

*Functional MRI studies* (20 studies). An ALE analysis based on only functional MRI studies produced 3 peak regions that mirrored areas observed in the overall and whole-brain only ALEs. Specifically, the functional MRI ALE included clusters in the right anterior insula (x = 26, y = 22, z = −4; 8 of 20 studies), the left anterior insula (x = −36, y = 16, z = −4; 8 of 20 studies) and the precuneus (x = 6, y =-68, z = 48; 5 of 20 studies); see Figure3A, Table 1, and Supplemental Table S4A for focus-count breakdowns for each cluster.

*Structural MRI studies* (9 studies). The structural MRI ALE produced only one consensus cluster, which was again located in the right anterior insula (x = 32, y = 16, z = −10) and showed relatively high reliability (4 of 9 studies) in the focus count analysis; see Figure3B, Table 1, and Supplemental Table S4B for a focus-count breakdown of this cluster.

*Social Media Use studies* (10 studies). An ALE based on the subset of 10 studies specifically investigating variation in social networking site use, which included 8 functional and 2 structural studies, produced a cluster that was again centered in the right anterior insula (x = 24, y =12, z =-8; Figure 3C, Table 1), with a peak slightly more medial to that found in the other ALE’s. As in the prior ALE maps, this cluster spanned an area that also included the inferior frontal gyrus and midline striatal regions. At a stricter cluster defining threshold of p <.001, we once again observed that only a circumscribed anterior insula cluster survived (see Supplemental Figure S1b). Despite the small number of studies included in this analysis, a focus-count analysis revealed moderate reliability across studies, with 5 of 10 papers contributing to this cluster (Supplemental Table S5B).

**Figure 3.**
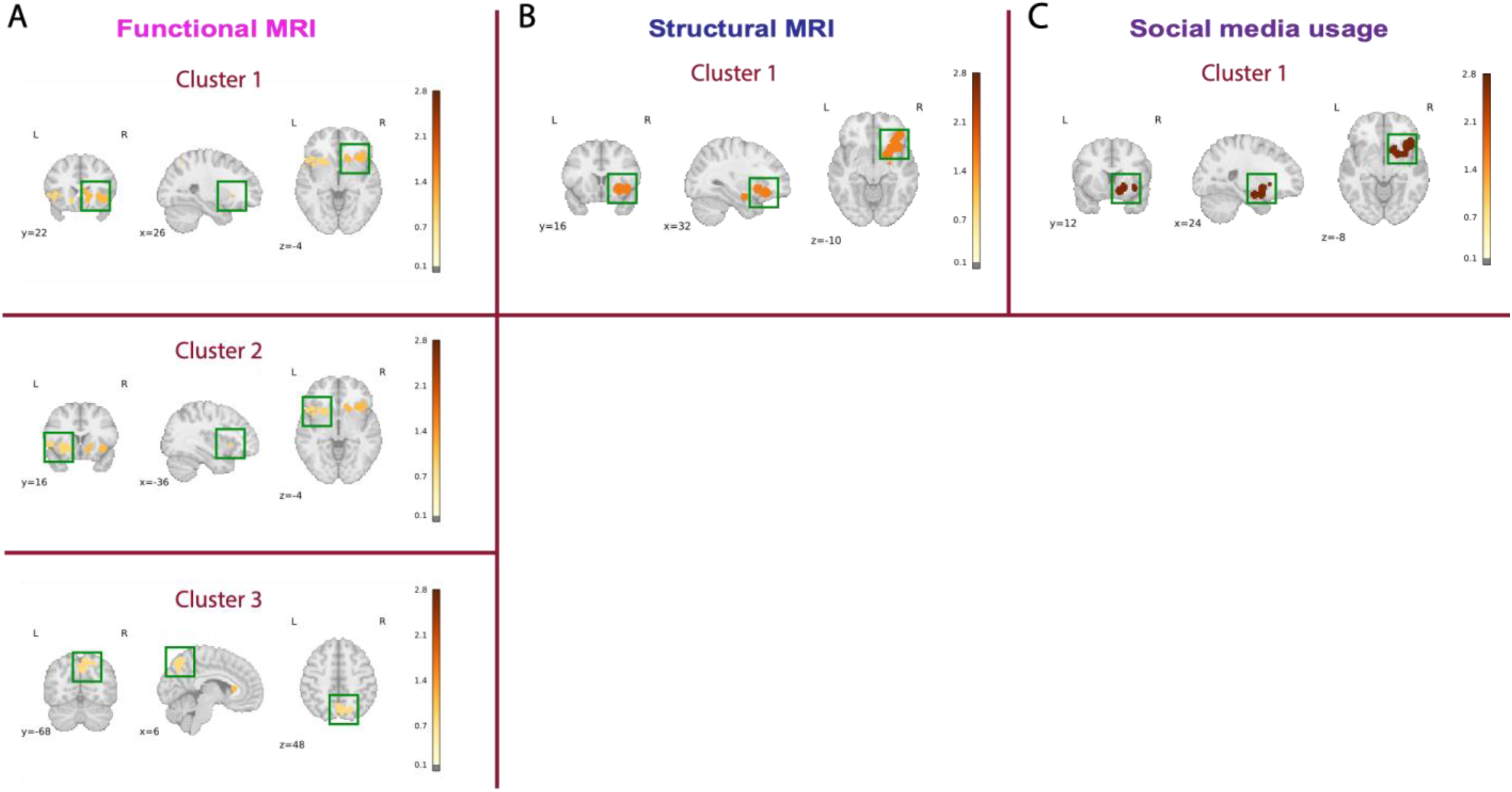
Individual ALEs separated by imaging modality and social media specificity. Three separate ALEs were run. The first two ALEs included separate studies based on imaging modality type: functional MRI and structural MRI. A third ALE included only studies that assessed social media use specifically, and included studies of both imaging modalities. A. The functional MRI ALE produced 3 peak clusters: two in the right and left anterior insula, and one encompassing the precuneus. B. The structural MRI ALE revealed one peak cluster again centered in the right anterior insula. C. The social media usage ALE also revealed one peak cluster in the right anterior insula, overlapping with clusters found in both the functional MRI and structural MRI ALE.

*Other forms of digital media behavior* (19 studies). For completeness, we also conducted a parallel analysis of the findings from studies that were not focused on social networking site use *per se*. This grouping of papers included a more heterogenous array of digital media behaviors (e.g., smartphone usage/dependency, overall screen-time measurements, internet usage/dependency). This collection of studies contributed to both the anterior insula and precuneus clusters that appear in the overall ALE (results described in Supplemental Material, and cluster focus counts reported in Supplemental Table S5A).

### Directionality of Effects

To understand the directionality of the individual study effects that contributed to significant clusters, we examined the frequency of positive and negative findings in each region (Table 1). We allowed for double counts in the case that a paper reported both negative and positive findings that contributed to the same cluster. Among the studies included in the ALE analyses, there were numerically more papers reporting negative (12 overall; 9 whole-brain) than positive (7 overall; 7 whole-brain) effects. Of note, each significant cluster in the overall ALE included a mix of positive and negative findings. The functional ALE clusters also reflected this mix of positive (11) and negative (14) effects, potentially signaling important nuances in the individual tasks and study designs, or in the way that digital media behavior was measured. There was considerably more consistency among the smaller number of papers that contributed to the structural ALE clusters, all of which reported effects in the negative direction (i.e., heavy digital media use was always reflected in decreased regional gray matter (cortical thickness and/or gray matter volume). The social ALE cluster centered in the anterior insula also had more negative (4) than positive (2) contributing findings and thus supported a trend for reduced activity/grey matter volume in association with heavier social media usage.

### Studies not included in ALE analyses

In an effort to capture the broader literature, we also conducted a region-count analysis based on effects reported in papers that passed all our inclusion criteria, but did not report MNI coordinates (n = 22). Supplemental Table S6 summarizes the regions where significant findings emerged in these papers. The most commonly reported effects in these region/network focused studies occurred in medial frontal (11/22) and lateral frontal (10/22) areas. While there is some very limited alignment between the specific loci reported in these studies and the clusters identified in our CBMA analyses — e.g. there are studies reporting significant effects in the anterior insula (n = 1) and precuneus (n = 1) — it is also clear that spatially targeted studies were more focused on brain regions not substantially represented in the ALE consensus maps (e.g., superior frontal gyrus, anterior cingulate cortex, amygdala).

## DISCUSSION

The central aim of this study was to objectively and quantitatively summarize findings in the growing neuroimaging literature focused on the brain correlates of habitual digital media engagement. Using a coordinate-based meta-analytic (ALE) approach, we find a small number of brain regions that show reliable spatial convergence across studies, despite cross-study differences in imaging modality, sample characteristics, and operationalizations of digital media use. Across analyses, the most consistently implicated region was the anterior insular cortex (bilaterally, but especially consistent in the right hemisphere), with less consistent involvement of the precuneus. Importantly, this pattern diverges in notable ways from the brain regions and networks most frequently emphasized in prior theoretical and empirical work, which has often focused on executive control systems, canonical reward circuitry, and the “social brain”.

### Anterior insula involvement across digital media behaviors

By far the most robust finding from the present analyses was the consistent involvement of the anterior insula. Peaks in this region were observed across the overall ALE, the whole-brain-only ALE, and ALEs restricted to functional MRI, structural MRI, and social media use studies. This convergence across imaging modalities and across different indices of digital media engagement suggests that anterior insula involvement likely reflects a general feature of digital media habits, rather than an effect tied to a particular task, analytic strategy, or behavioral measure. Despite this relative consistency in our study, the anterior insula has received relatively little emphasis in prior attempts to summarize the digital media neuroimaging literature (cf., Maza et al., 2023; Wadsley & Ihssen, 2023). Indeed, among the 22 studies that met inclusion criteria but eschewed whole-brain stereotaxic reporting in favor of a focus on specific regional findings, only one reported an effect proximal to the anterior insula (which the authors localized to the inferior frontal gyrus). This clear discrepancy underscores the value of using a data-driven meta-analytic approach to identify neural loci that may be underrepresented in theory-guided reviews or region-focused studies.

Of course, it should also be acknowledged that, despite being the most consistently implicated area, the anterior insula appeared in fewer than half of the overall studies included in the ALE (13 of 29) and effects were not always in the same direction (5 findings were in the “positive” direction and 9 in the “negative direction). Accordingly, we should be careful not to characterize findings in this region as a ubiquitous neural signature of digital media use behavior. Rather, the regular emergence of the anterior insula across studies suggests that it represents a reliable, though not universal, locus of variation associated with digital media habits.

The anterior insula has most often been highlighted in the literature for its role in the integration of cognitive and socio-emotional processing (Uddin et al., 2017) and in salience detection (along with the anterior cingulate; Menon & Uddin, 2010). It is frequently engaged during anticipation of potentially significant or threatening events (Meyer et al., 2019), in heightened anxiety states (Klumpp et al., 2012; Simmons et al., 2011), and in response to social exclusion and rejection3/5/2026 8:21:00 PM(Eisenberger et al., 2003; Masten et al., 2009) –– processes that are relevant to many digital media contexts, particularly social media environments that emphasize social evaluation, novelty, and uncertainty. At the same time, the anterior insula is increasingly understood as a region that integrates salient sensory, affective, and interoceptive information with ongoing cognition in order to guide behavior and action (Dosenbach et al., 2025). Through connections with the dorsal anterior cingulate cortex, lateral prefrontal cortex, and premotor regions, the anterior insula is well positioned to influence downstream control and action systems. From this integrative perspective, anterior insula involvement in digital media use may reflect group and individual differences in how salient digital cues are registered and translated into engagement-related action. In this way, variation in anterior insula structure and function may index differences in how strongly or efficiently salient cues, such as notifications, social feedback, or novel content, trigger shifts toward externally oriented, action-ready, states. This interpretation also accommodates findings linking the anterior insula to craving and habitual behavior across multiple domains, including substance use (Montag & Becker, 2023) and problematic technology use (Turel et al., 2021).

### Precuneus involvement and memory processing

The precuneus appeared as a weaker, yet recurring, consensus region in the overall ALE and in ALEs based on functional MRI and heterogeneous digital media measures. As a central hub of the default mode network (Utevsky et al., 2014), the precuneus is commonly associated with processes that are more strongly activated when we are at rest (Greicius et al., 2003; Zhang & Li, 2012) in order to enact internally oriented cognitive processes, including episodic memory retrieval (Foudil & Macaluso, 2024; Jonker et al., 2018), mental imagery (Fletcher et al., 1995; Lundstrom et al., 2003), and self-referential processing (Herold et al., 2016; Sajonz et al., 2010). Its’ absence from the structural MRI ALE, and the relatively modest focus counts obtained in this region, suggest that precuneal involvement marks state-dependent patterns of engagement rather than stable trait-level differences. One possibility is that habitual digital media use influences internally oriented processing during passive content consumption, such as scrolling or media viewing, which may share features with movie-watching paradigms known to engage the precuneus (Cooper et al., 2021; Demirtaş et al., 2019). Alternatively, precuneus involvement may reflect memory-related processes, such as event segmentation (Baldassano et al., 2017; Speer et al., 2007) or incidental encoding (Foudil et al., 2020; Kauttonen et al., 2018), that occur during extended digital media exposure. Although speculative, these interpretations point to potential mechanisms that have received little attention in the current literature and may warrant further investigation.

## Limitations

Given that this domain of research is relatively new, and has thus far produced a still modestly sized literature, we took a broad approach to study inclusion. In doing so, we included studies using multiple imaging modalities, varied age groups, and disparate indices of digital media behavior. While taking this inclusive approach maximized the number of studies that could be incorporated into the ALEs, it also limits the findings to effects that are tolerant to cross-study heterogeneity. For instance, the inclusion of studies from both developmental samples (children and adolescents) increased the number of studies contributing to each ALE, but may have veiled the involvement of regions that experience changes in the relationship to digital media habits across development (e.g., age differences in the structure and/or function of the prefrontal cortex might dilute consensus effects in that region in the adult age group). Currently, there are too few studies within narrow age-bands to run separate age-group specific ALEs. Indeed, current best practices in CBMA meta-analysis recommend at least 17-20 studies for each ALE (Müller et al., 2018), and while the overall study ALE included 29 studies, some ALEs using subsets of studies fell below this recommended minimum (e.g., there were only 9 studies contributing to the structural MRI ALE). The limited number of available studies supporting these subset ALE’s thus urges cautious interpretation of the findings they produced. Further interpretive caution is merited by the fact that the significant ALE clusters also typically reflected coordinates from a mixture of positive (greater activation/gray matter) and negative (reduced activation/gray matter) effects.

### Conclusions and Implications for models of digital media engagement

This study represents, to our knowledge, the first quantitative coordinate-based meta-analysis of neuroimaging studies examining habitual digital media engagement. Using a data-driven approach, we identified consistent convergence in a small set of brain regions, most prominently the anterior insula (extending toward the inferior frontal gyrus and ventral striatum) and the precuneus. Taken together, the present findings suggest that habitual digital media engagement may be most consistently associated with the neural system involved in evaluating salience, integrating socio-emotional relevance, and translating such information into engagement-related behavior, rather than with executive control, reward processing, or social information processing in isolation. The relative absence of consistent prefrontal convergence in this meta-analysis challenges models that frame digital media use primarily in terms of top-down control deficits (e.g. Chiu & Chein, 2022; Crone & Konijn, 2018; Hutton et al., 2024; Meshi et al., 2015).

Likewise, the inconsistency of findings in the striatum and precuneus should temper over-emphasis on reward- and social information processing frameworks. Instead, the most reliable neural locus across studies was in the anterior insula, potentially acting as a bottleneck through which salient digital cues influence engagement. By highlighting this region, the present meta-analysis refines our understanding of the brain systems most reliably associated with digital media habits and orients us to a relatively under-emphasized target region for further longitudinal, mechanistic, and intervention studies.

## Notes

**Declarations**: This work was supported in part by a grant from the Eunice Kennedy Shriver National Institute of Child Health and Human Development to J. Chein, R01 HD098097. The authors have no relevant financial or non-financial interests to disclose. Approval was obtained from an IRB committee of Temple University and adhere to the tenets of the Declaration of Helsinki. The datasets generated during and/or analyzed in the current study are available from the corresponding author upon reasonable request.

### Competing Interest Statement

The authors have declared no competing interest.

