## Supplemental Files for "Habitual digital media use and the brain: a meta-analysis"

**SUPPLEMENTAL TABLES AND ANALYSES**

**Supplemental Figure S1.**

**Overall ALE**

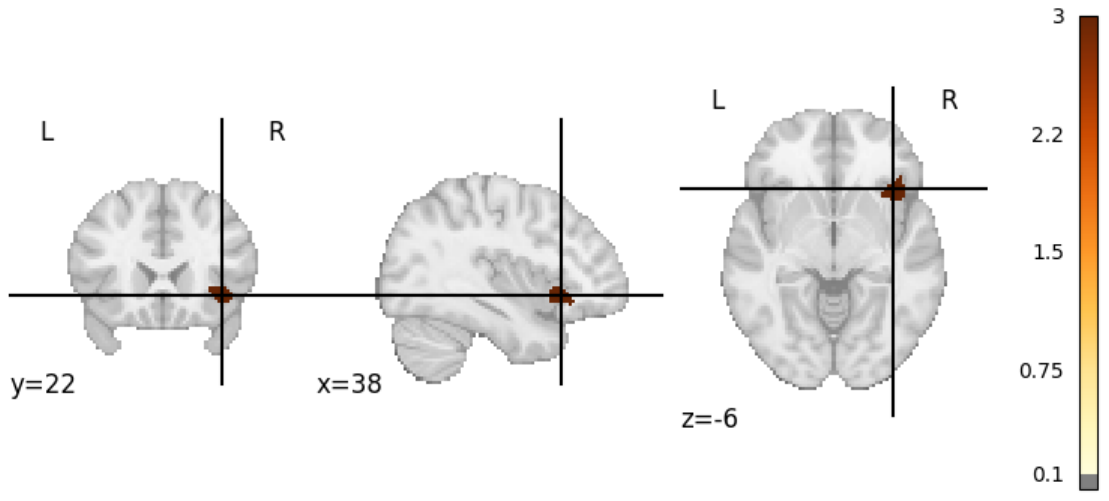

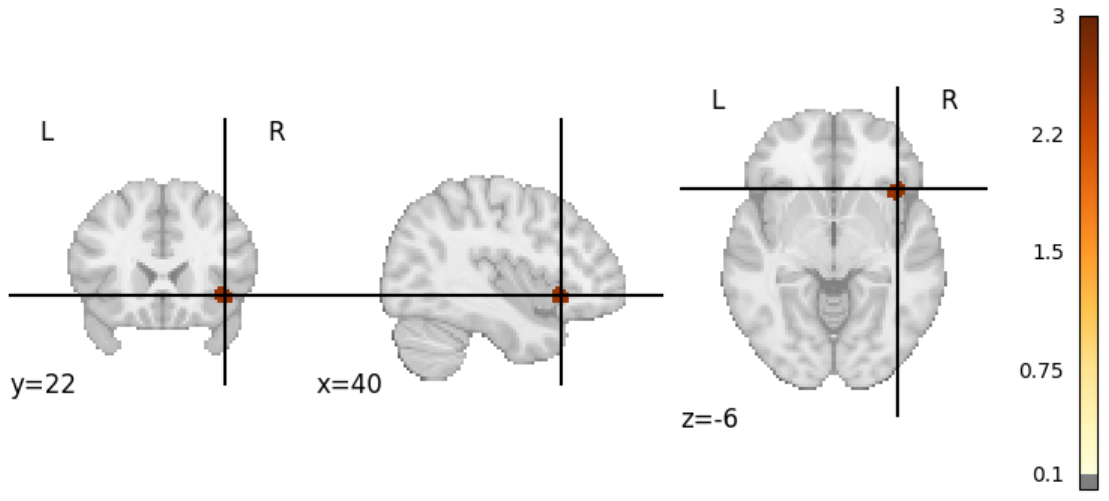
**ALE of Social Media Studies**

**Figure S1. Anterior insula clusters identified with cluster defining threshold set a p-value of p < .001 for the (top) overall ALE and the (bottom) social media studies ALE. The crosshairs are centered on the peak location within each cluster.**

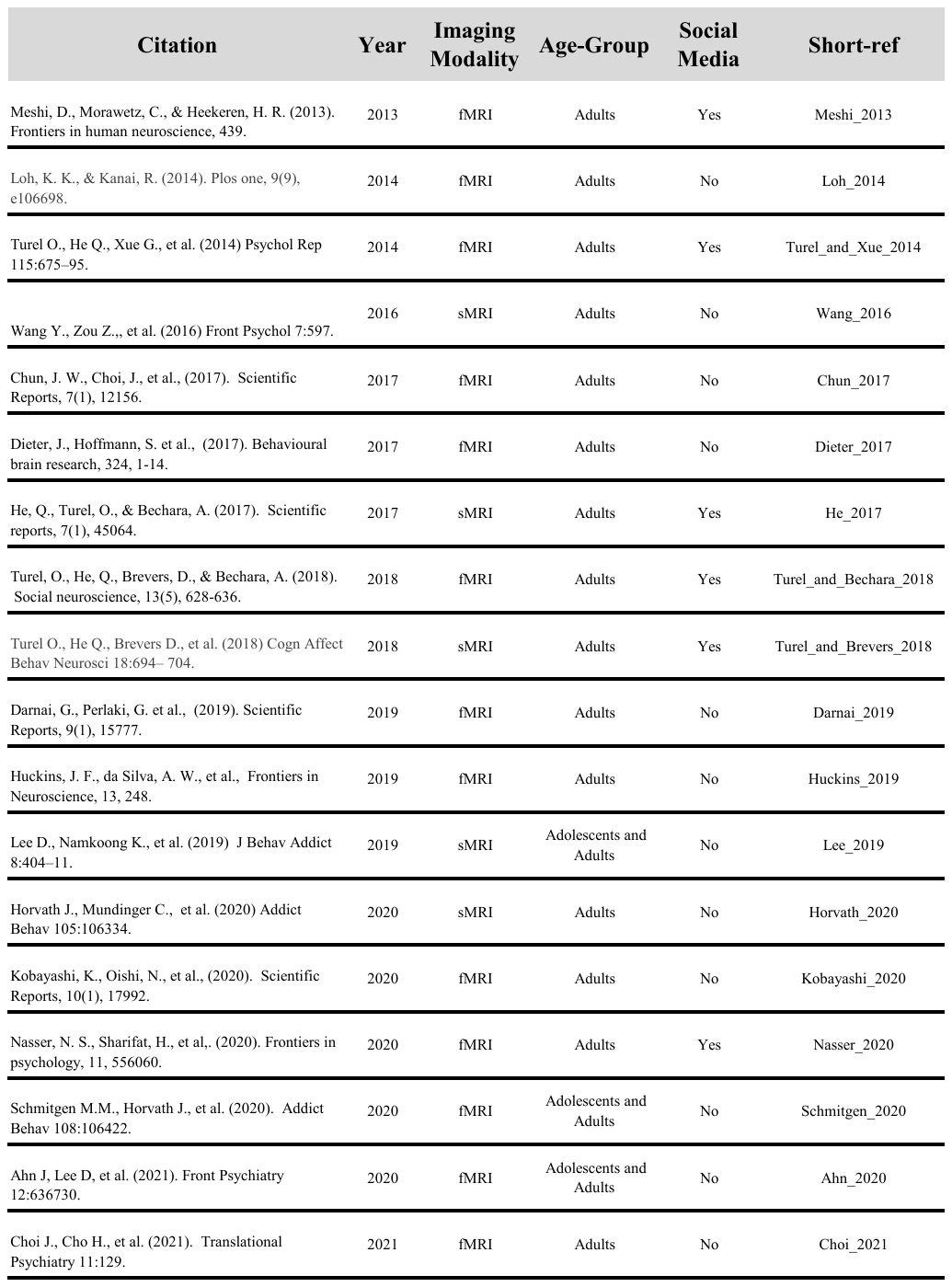
**Supplemental Table S1. All studies included in Overall ALE (n = 29)**

**Supplemental Tables S2A-F: Peak Clusters for ALEs**

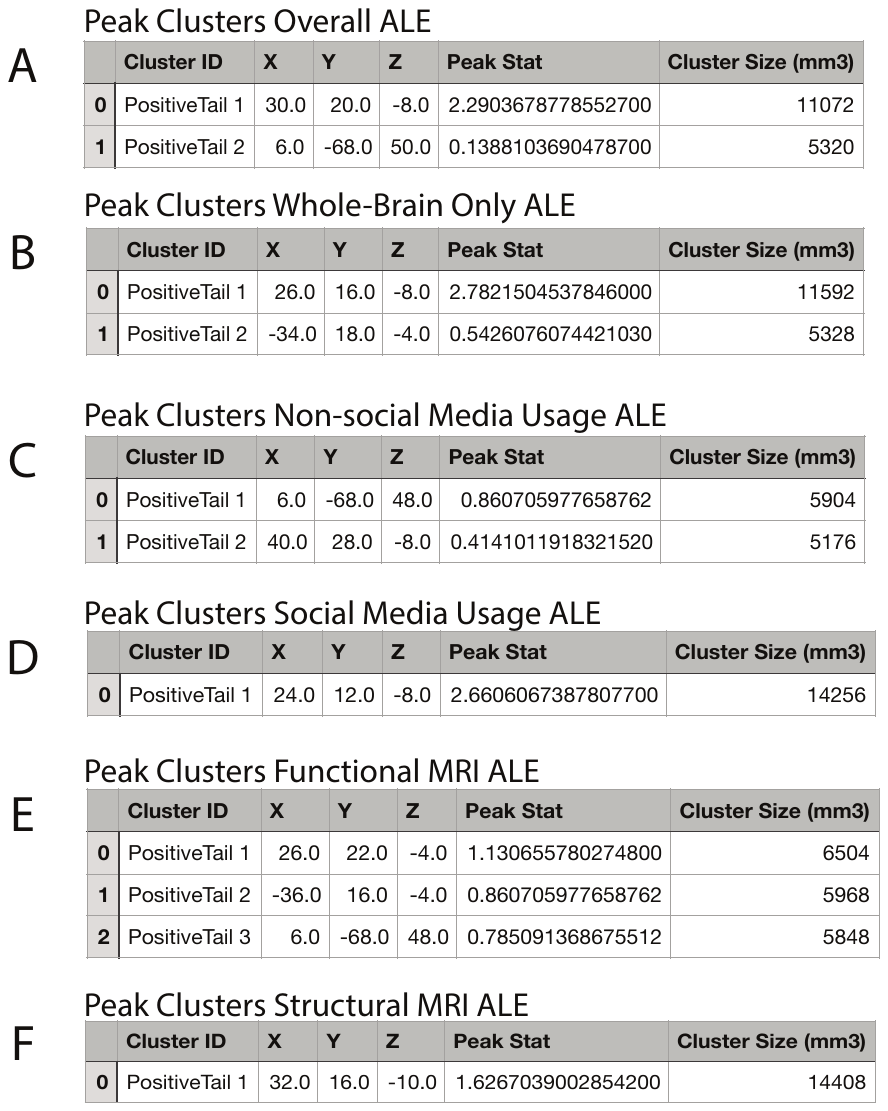

**Supplemental Table S3A/B: Focus Count Results for Overall and Whole Brain ALEs**

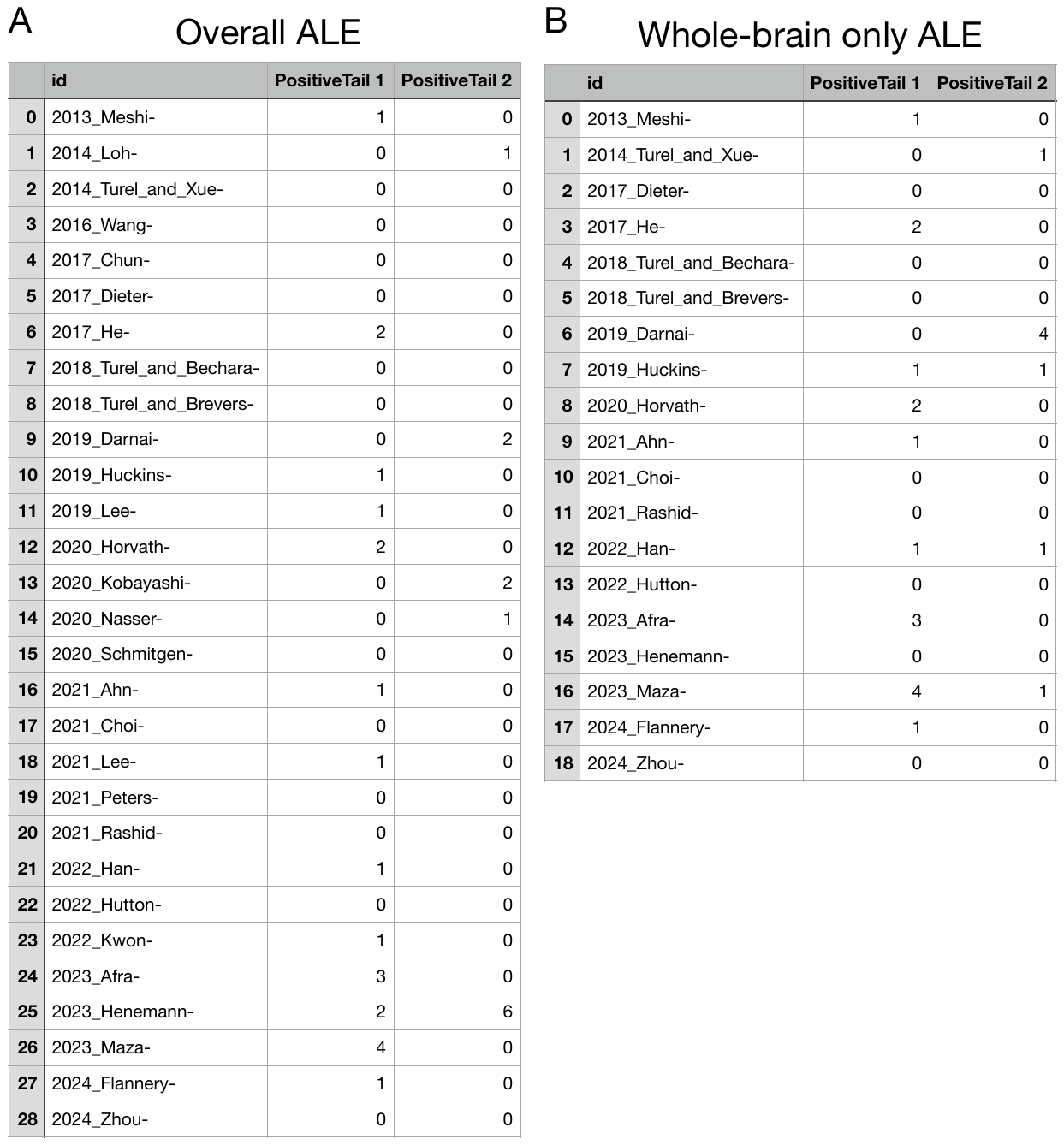

**Supplemental Table S4A/B Focus Count Results for Imaging Modality ALEs**

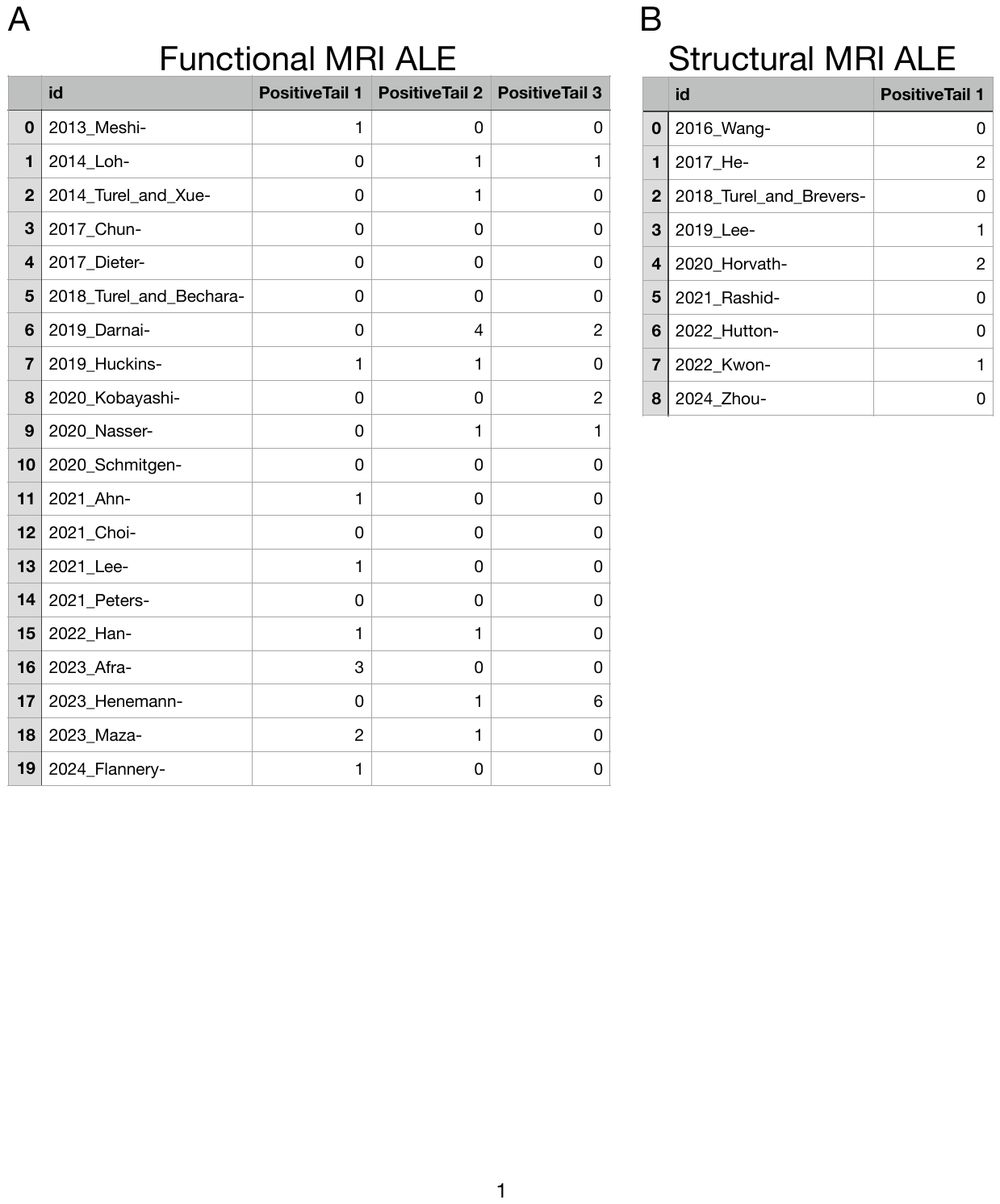

**Supplemental Table S5A/B Focus Count Results for Usage Type ALEs**

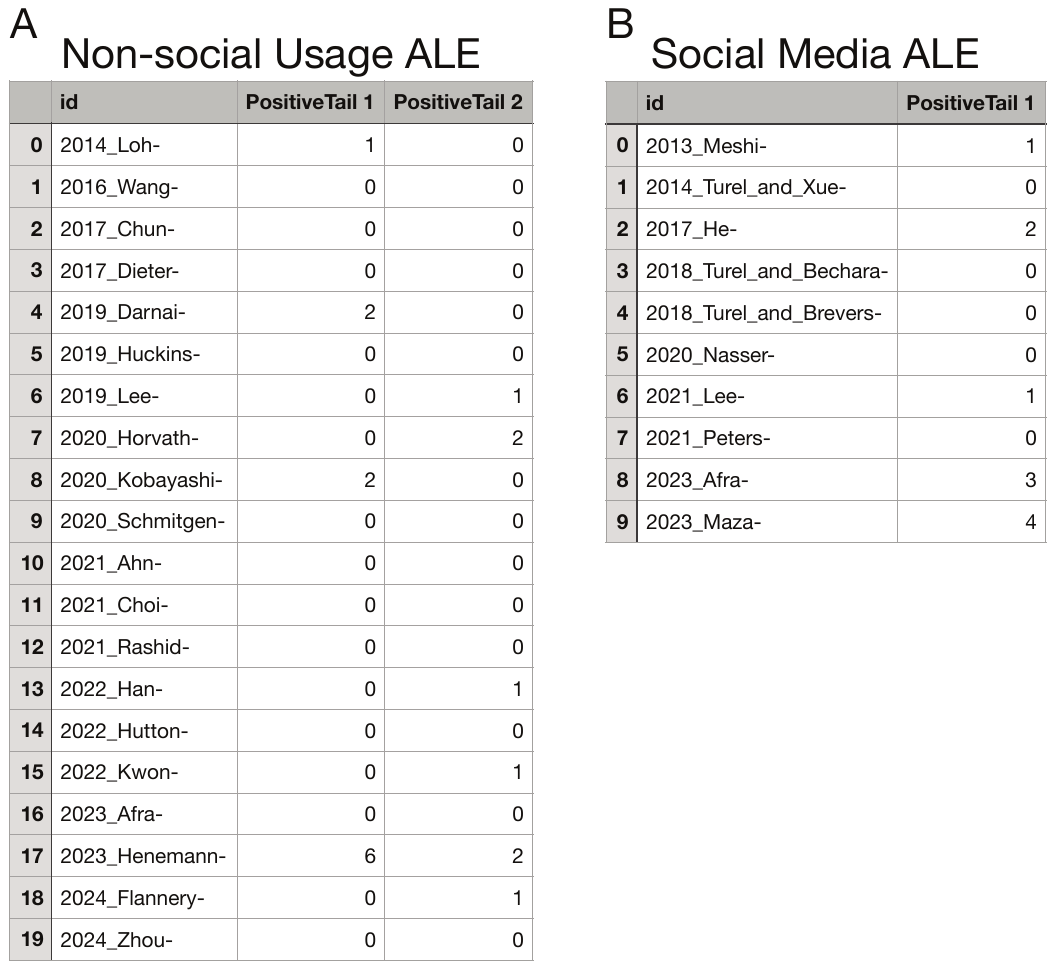

**“Other” digital media behavior subset ALE**

*Studies of other digital media behaviors* (19 studies). Two significant clusters emerged from the set of 19 studies evaluating digital use habits other than social media use. Peak one (x = 6, y = -68, z = 48) aligned with the precuneus peak obtained in the overall ALE. Although, as in the overall ALE, a focus-count analysis revealed that this peak was only supported by 4 of 20 studies with an outsize contribution from Henemann et al., 2023 (see Supplemental Table 4A). Peak two (x = 40, y = 28, z = -8) was centered in the right inferior frontal gyrus but extended into the right anterior insula region also identified in the overall ALE (see Supplemental Table 2C for information on both peaks). Focus-count analyses revealed that this second peak had modest reliability across studies (6 of 20).

**Findings from ROI and Network-based Studies**

Since these papers used varied functional and structural labels to describe regions where effects were observed, we reviewed (through visual assessment and cross-referencing to commonly used brain atlases and Neurosynth) and re-categorized significant findings as closely and specifically as possible, and according to the coarse taxonomy of regions seen in Table S6. As with the CBMA, we also coded studies according to imaging modality and directionality of effects, and then tallied the total number of significant effects reported in each major region category, across the 22 papers.

Lastly, a region that was only reported as a significant region once (1/22) in this qualitative count, but features prominently in our ALE peaks is the precuneus. This may indicate that the precuneus is noteworthy in that it is not often identified as an a-priori region of interest. However, the precuneal peaks were less robust and inconsistent across the individual ALEs compared to the anterior insula peaks so it could be that this activation is simply not as reliable.

Overall, we find some consistency between our CBMA in these relevant papers that conducted apriori ROI-based analyses – particularly with the prevalence of regions located in Lateral Frontal and subcortical regions. However, despite the prevalence of medial frontal regions like the mPFC in this qualitative analysis and the literature at large, we do not see this in our CBMA. We also see modest evidence of a precuneal effect in our CBMA that only appeared once in this qualitative analysis and is rarely reported in the existing literature.

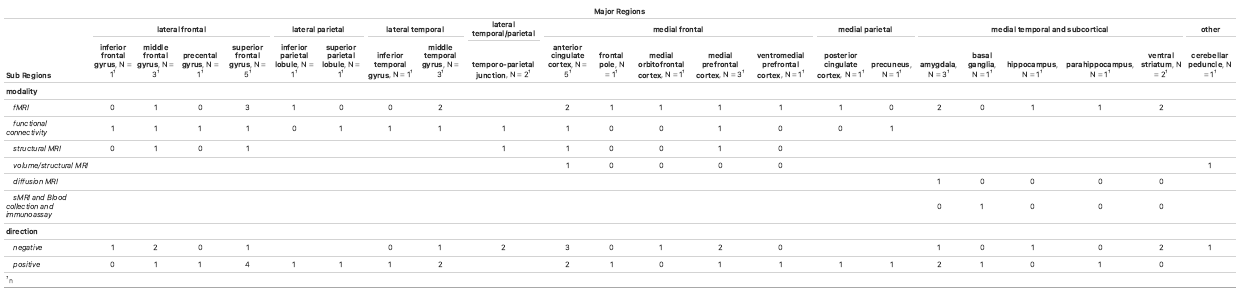
Supplemental Table S6
